# Extended Modelling of Platelet Calcium Signaling by Combined Recurrent Neural Network and Partial Least Squares Analyses

**DOI:** 10.1101/2025.02.01.636037

**Authors:** Chukiat Tantiwong, Hilaire Yam Fung Cheung, Joanne L Dunster, Jonathan M. Gibbins, Johan W. M. Heemskerk, Rachel Cavill

## Abstract

Platelets play critical roles in hemostasis and thrombosis. The platelet activation process is driven by agonist-induced rises in cytosolic [Ca^2+^]_i_, where the patterns of Ca^2+^ responses are still incompletely understood. In this study, we developed a number of techniques to model the [Ca^2+^]_i_ curves of platelets from a single blood donor. Using a fluorescence ratio probe, the platelets were stimulated with a panel of agonists, i.e. thrombin, collagen, or CRP under various conditions, preventing extracellular Ca^2+^ entry, secondary mediator effects or Ca^2+^ reuptake into intracellular stores. To analyze the data, we developed two non-linear models, a multilayer perceptron (MLP) network and an autoregressive network with exogenous inputs (NARX). The trained networks accurately predicted the platelet [Ca^2+^]_i_ curves in the presence of combinations of agonists and inhibitors, with the NARX model achieving an R^2^ up to 0.64 for trend prediction of unforeseen data. In addition, we used the same dataset for construction of a partial least square (PLS) linear regression model, which estimated the explained variance of each input. The NARX model demonstrated that good fits could be obtained for the calcium curves modelled, whereas the PLS model gave useful interpretable information on the importance of each variable. These modelling results can be used for the development of novel platelet [Ca^2+^]_i_-inhibiting drugs.

## 1. Introduction

Blood platelets, derived from megakaryocytes, function in hemostasis and thrombosis via receptor-induced signaling responses [1–3]. Important platelet-activating receptors are the protease-activated receptors (PAR1/4) for thrombin and the glycoprotein VI (GPVI) receptor for collagen, which signal as G-protein coupled receptors (GPCR) and as a protein tyrosine kinase-linked receptor (TKLR), respectively [4]. Given that arterial thrombosis is driven by the activation and aggregation of platelets [5], and it is a prominent cause of death worldwide [6], a clear understanding of the process of platelet activation is a must.

In platelets stimulated via GPCR or TKLR, a rise in cytosolic [Ca^2+^]_i_ is the common initial event, mediating all essential platelet functions [7,8]. The agonist-induced mobilization of Ca^2+^ from intracellular stores in the endoplasmic reticulum (or dense tubular system) occurs via inositol 1,4,5-trisphosphate receptors (IP_3_Rs), whereas sarcoplasmic/endoplasmic reticulum Ca^2+^-ATPases (SERCAs) are responsible for Ca^2+^ back pumping into these stores (Figure S1) [7,8]. The IP_3_R channels are triggered by IP_3_, which is produced as a result of activation of the GPCRs for thrombin [9] and ADP [10], and of activation of the TKLR GPVI by collagen or collagen-related peptide (CRP) [8].

According to the mechanism of store-operated Ca^2+^ entry (SOCE), the store depletion is coupled to entry of Ca^2+^ from the extracellular medium, via Orai1 channels, which then interact with a Ca^2+^ sensor STIM1 (stromal interaction molecule 1) in the endoplasmic reticulum membrane [7]. The back pumping of Ca^2+^ over the plasma membrane occurs via plasma membrane Ca^2+^-ATPases (PMCAs). The primary agonists, thrombin and CRP, furthermore stimulate the release of autocrine agents that can enforce the Ca^2+^ signaling process. These are in particular autocrine-produced thromboxane A_2_ (TxA_2_) and ADP, both of which stimulate IP_3_ production via GPCRs [11]. Another paracrine-dependent Ca^2+^ entry mechanism is provided by ATP, which activates P2X_1_ channels that specifically mediate Ca^2+^ entry [12].

Several pharmacological inhibitors can be used to interfere with the platelet Ca^2+^ responses. The entry of Ca^2+^ from the extracellular fluid is prevented by the Ca^2+^ chelator EGTA. Back pumping of Ca^2+^ from cytosol to intracellular stores is inhibited by the compound thapsigargin, which accordingly potentiates the Orai1-STIM1 dependent entry [7]. The effects of autocrine agents can be suppressed by the presence of apyrase (degrading ATP and ADP) and indomethacin (blocking TxA_2_ formation). Figure S1 illustrates the platelet receptors, ligands, inhibitors and channels, relevant to the present study.

The high complexity of the Ca^2+^-modulating process in platelets has triggered other authors to develop mathematical models, aiming not only to better understand the process but also to identify new therapeutic targets. Other authors combined several models of Ca^2+^ fluxes in different platelet compartments into one single system, using a set of ordinary differential equations (ODEs) [13]. Although their system did not include ligand-receptor interactions, it still consisted of 34 entities, 35 interactions and 86 parameters, thus reflecting the complexity of Ca^2+^ signaling process. An alternative approach presented by Chatterjee and Diamond [14] was to create a neural network model, which was trained from the Ca^2+^ response patterns to specific agonists, using the platelets from a number of healthy donors. The neural network, acting as a black box, could predict synergistic effects on the Ca^2+^ responses of up to six receptor agonists. A trade-off of this network model was that all the parameters needed to be tuned and trained, which required extensive experimental data to achieve the desired predictive power. Another limitation was that the neural network approach did not provide information on the contribution of each type of Ca^2+^ channel and pump to the overall [Ca^2+^]_i_ levels. Similarly, it did not identify how the blockage of a given channel or (autocrine) process influenced the overall response.

In the present study, we constructed a computational model to predict the shapes of platelet [Ca^2+^]_i_ curves over time in response to thrombin or CRP for a given set of experimental conditions, with known agonists and inhibitors. We built several neural network-based models to better predict the agonist and inhibitor effects on the [Ca^2+^]_i_ time curves. We subsequently used a partial least square regression analysis to understand how specific curve variables contributed to obtained response. To exclude inter-individual variation, we used a coherent set of Ca^2+^ response curves in platelets, taken from one representative experiment using platelets from a healthy subject.

## 2. Results

### 2.1. Comparing all Input Agonist-Induced Platelet [Ca ^2+^]_i_ Curves

Using a high-throughput method described before [15], the Fura-2-loaded platelets from a single healthy donor were incubated in the presence of EGTA or CaCl_2_ with or without secondary mediator inhibitors apyrase and indomethacin (AI); and then stimulated with collagen, CRP or thrombin. Under all these conditions, agonist-induced rises in [Ca^2+^]_i_ were measured as nM concentrations over a time period of 9 min. Altogether, by also varying the agonist concentrations, resulted in a set 72 different experimental conditions (Table 1). For the present paper, the 72 different [Ca^2+^]_i_ time curves were selected of the platelets from one donor, which were most representative for the set of 6 donors [15].

**Table 1.** Assignment matrix of variables of experimental conditions. Fura-2-loaded platelets from one single donor were used for Ca^2+^ response measurements on the same day. Conditions highlighted in blue (solid borders) were used as validation set, while those in red (dashed borders) were used as test set. *Abbreviations:* No., condition number; Col, collagen (μg/mL); CRP, collagen-related peptide (μg/mL); Thr, thrombin (nM); EG, EGTA: 0.1 mM if assigned to 1; or 1 mM CaCl_2_ if assigned to 0; AI, apyrase (0.1 U/mL) plus indomethacin (20 μM); Thap, thapsigargin (1 μM).

| No. | Col | CRP | Thr | EG | AI | Thap |
| --- | --- | --- | --- | --- | --- | --- |
| 1 | 0 | 0 | 0 | 0 | 1 | 0 |
| 2 | 0 | 0 | 0 | 1 | 1 | 0 |
| 3 | 0 | 0 | 0 | 0 | 1 | 1 |
| 4 | 0 | 0 | 0 | 1 | 1 | 1 |
| 5 | 0 | 0 | 0 | 0 | 0 | 0 |
| 6 | 0 | 0 | 0 | 1 | 0 | 0 |
| 7 | 0 | 0 | 0 | 0 | 0 | 1 |
| 8 | 0 | 0 | 0 | 1 | 0 | 1 |
| 9 | 30 | 0 | 0 | 0 | 1 | 0 |
| 10 | 10 | 0 | 0 | 0 | 1 | 0 |
| 11 | 3 | 0 | 0 | 0 | 1 | 0 |
| 12 | 1 | 0 | 0 | 0 | 1 | 0 |
| 13 | 30 | 0 | 0 | 1 | 1 | 0 |
| 14 | 10 | 0 | 0 | 1 | 1 | 0 |
| 15 | 3 | 0 | 0 | 1 | 1 | 0 |
| 16 | 1 | 0 | 0 | 1 | 1 | 0 |
| 17 | 30 | 0 | 0 | 0 | 1 | 1 |
| 18 | 10 | 0 | 0 | 0 | 1 | 1 |
| 19 | 3 | 0 | 0 | 0 | 1 | 1 |
| 20 | 1 | 0 | 0 | 0 | 1 | 1 |
| 21 | 30 | 0 | 0 | 1 | 1 | 1 |
| 22 | 10 | 0 | 0 | 1 | 1 | 1 |
| 23 | 3 | 0 | 0 | 1 | 1 | 1 |
| 24 | 1 | 0 | 0 | 1 | 1 | 1 |
| 25 | 10 | 0 | 0 | 0 | 0 | 0 |
| 26 | 1 | 0 | 0 | 0 | 0 | 0 |
| 27 | 10 | 0 | 0 | 1 | 0 | 0 |
| 28 | 1 | 0 | 0 | 1 | 0 | 0 |
| 29 | 10 | 0 | 0 | 0 | 0 | 1 |
| 30 | 1 | 0 | 0 | 0 | 0 | 1 |
| 31 | 10 | 0 | 0 | 1 | 0 | 1 |
| 32 | 1 | 0 | 0 | 1 | 0 | 1 |
| 33 | 0 | 10 | 0 | 0 | 0 | 0 |
| 34 | 0 | 1 | 0 | 0 | 0 | 0 |
| 35 | 0 | 10 | 0 | 1 | 0 | 0 |
| 36 | 0 | 1 | 0 | 1 | 0 | 0 |
| No. | Col | CRP | Thr | EG | AI | Thap |
| 37 | 0 | 10 | 0 | 0 | 0 | 1 |
| 38 | 0 | 1 | 0 | 0 | 0 | 1 |
| 39 | 0 | 10 | 0 | 1 | 0 | 1 |
| 40 | 0 | 1 | 0 | 1 | 0 | 1 |
| 41 | 0 | 10 | 0 | 0 | 1 | 0 |
| 42 | 0 | 1 | 0 | 0 | 1 | 0 |
| 43 | 0 | 10 | 0 | 1 | 1 | 0 |
| 44 | 0 | 1 | 0 | 1 | 1 | 0 |
| 45 | 0 | 10 | 0 | 0 | 1 | 1 |
| 46 | 0 | 1 | 0 | 0 | 1 | 1 |
| 47 | 0 | 10 | 0 | 1 | 1 | 1 |
| 48 | 0 | 1 | 0 | 1 | 1 | 1 |
| 49 | 0 | 0 | 10 | 0 | 1 | 1 |
| 50 | 0 | 0 | 3 | 0 | 1 | 1 |
| 51 | 0 | 0 | 1 | 0 | 1 | 1 |
| 52 | 0 | 0 | 0.3 | 0 | 1 | 1 |
| 53 | 0 | 0 | 10 | 1 | 1 | 1 |
| 54 | 0 | 0 | 3 | 1 | 1 | 1 |
| 55 | 0 | 0 | 1 | 1 | 1 | 1 |
| 56 | 0 | 0 | 0.3 | 1 | 1 | 1 |
| 57 | 0 | 0 | 10 | 0 | 1 | 0 |
| 58 | 0 | 0 | 3 | 0 | 1 | 0 |
| 59 | 0 | 0 | 1 | 0 | 1 | 0 |
| 60 | 0 | 0 | 0.3 | 0 | 1 | 0 |
| 61 | 0 | 0 | 10 | 1 | 1 | 0 |
| 62 | 0 | 0 | 3 | 1 | 1 | 0 |
| 63 | 0 | 0 | 1 | 1 | 1 | 0 |
| 64 | 0 | 0 | 0.3 | 1 | 1 | 0 |
| 65 | 0 | 0 | 10 | 0 | 0 | 1 |
| 66 | 0 | 0 | 1 | 0 | 0 | 1 |
| 67 | 0 | 0 | 10 | 1 | 0 | 1 |
| 68 | 0 | 0 | 1 | 1 | 0 | 1 |
| 69 | 0 | 0 | 10 | 0 | 0 | 0 |
| 70 | 0 | 0 | 1 | 0 | 0 | 0 |
| 71 | 0 | 0 | 10 | 1 | 0 | 0 |
| 72 | 0 | 0 | 1 | 1 | 0 | 0 |

Comparing the set of original traces (Figure S2), several characteristics can be observed, except for the expected agonist dose-dependency [15]. In general, the [Ca^2+^]_i_ curves induced by the weak GPVI agonist collagen showed steady increases with lower maximal amplitudes (Exp. 9-31), when compared to the higher amplitude and often biphasic [Ca^2+^]_i_ rises induced by the strong GPVI agonist CRP (Exp. 37-48). In particular, the curves with the PAR1/4 agonist thrombin (Exp. 49-72) had a transient shape, indicating high activity of the SERCA Ca^2+^ pumps. Other differences were 4-80 times higher amplitude curves (depending on other variables) in the presence of CaCl_2_ than with EGTA, which in part was due to Orai1-dependent Ca^2+^ entry [15]. Furthermore, we observed potent [Ca^2+^]_i_ increase by adding the SERCA inhibitor thapsigargin, inhibiting SOCE and activating the Orai1 channels [7]. Effects of the autocrine inhibitors indomethacin and apyrase (IA) were a consistent lowering of most of the curves.

### 2.2. Workflow of the Modelling Approaches

In order to prepare the experimental data for further processing, we first interpolated and smoothened the 72 curves at 1 s time intervals (Figure S3), followed by a *y*-axis scaling per curve from 0-1 (Figure S4). The subsequent workflow (Figure 3) consisted of feature generation by combining and squaring of the experimental variables (see below), and split the curves into training, validation and test sets. The data were used as input for two types of modelling, i.e. neural network and PLS analyses. In the neural network analysis, we used the NARX procedure for trend prediction and the MLP procedure for magnitude prediction. A combined optimized network was tested on final performance. On the other hand, PLS was used to directly model the scalar characteristics of the curves. The result from both approaches was interpreted and cross-checked with each other.

**Figure 1.**
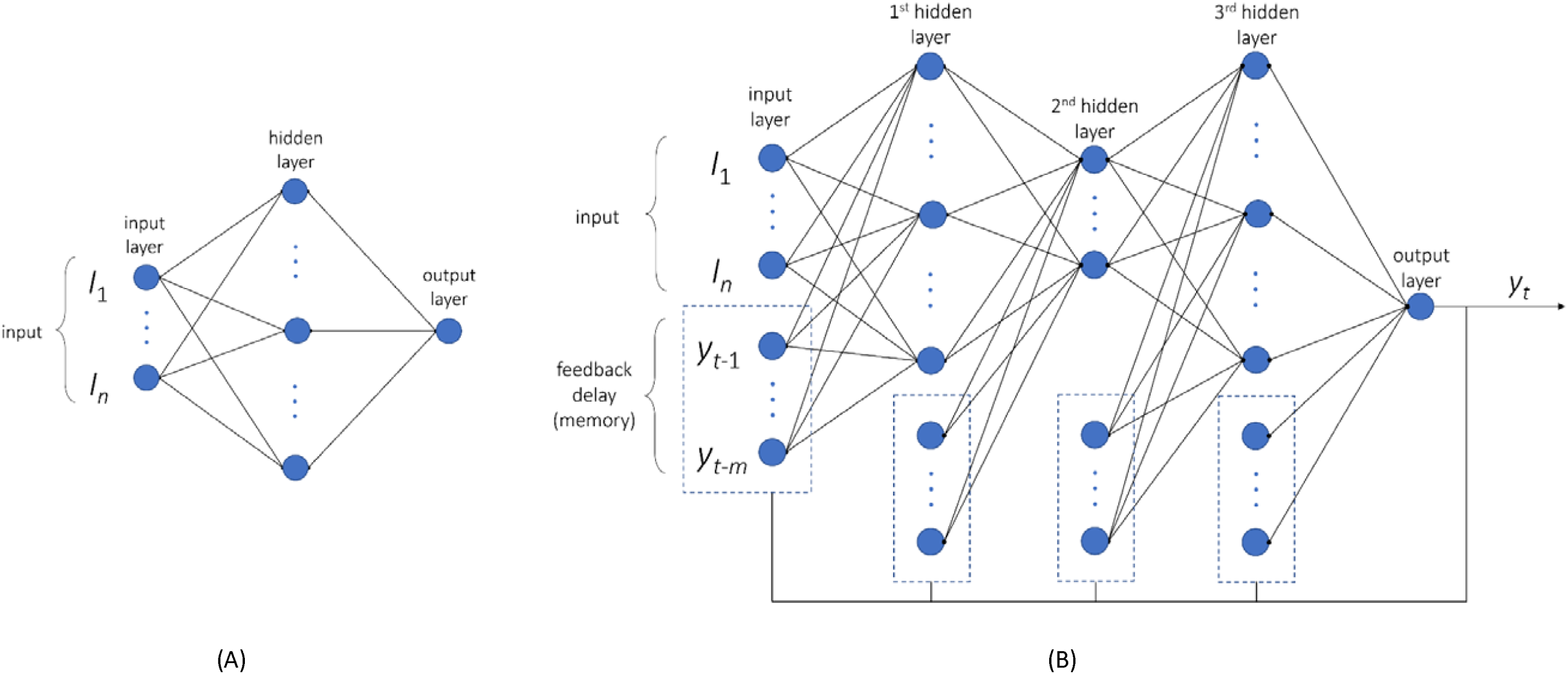
Construction of two neural networks. (**A**) Setup of MLP network, as a fully connected feedforward neural network, which was used for prediction of the magnitude of [Ca^2+^]_i_ time curves. (**B**) Closed-loop non-linear autoregressive network with exogenous inputs (NARX), which was used as a recurrent neural network. Herein, generated output served as input for a next time point. This network type was used for the prediction of trends in [Ca^2+^]_i_ time curves.

**Figure 2.**
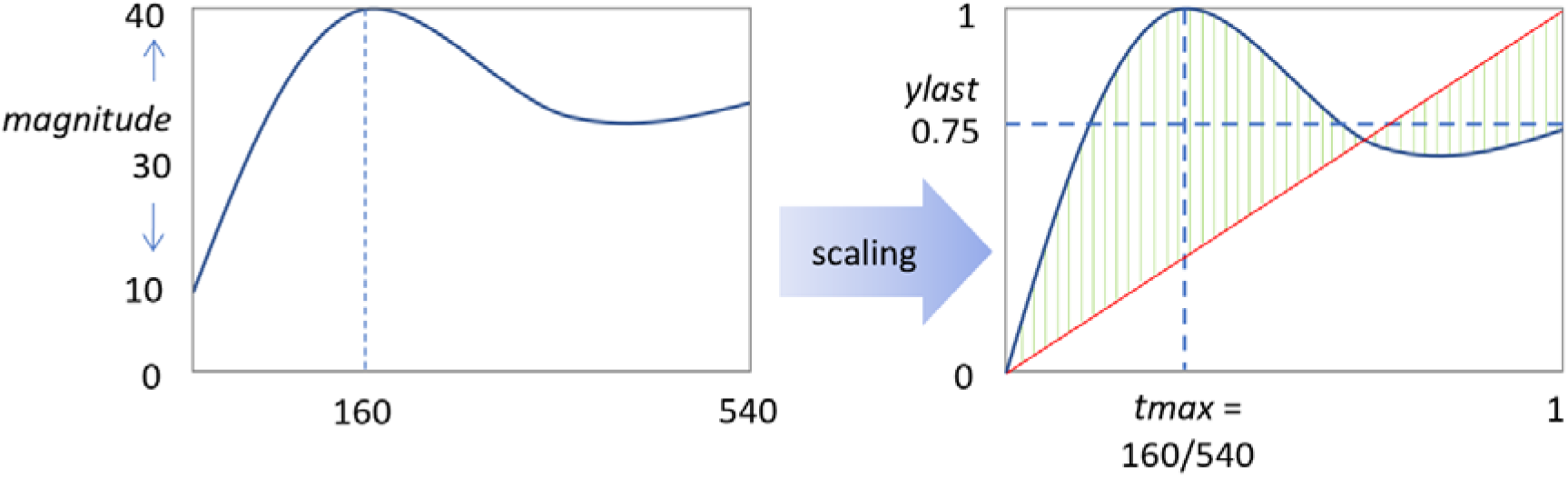
Defining the scalar characteristic of a [Ca^2+^]_i_ time curve. Scaling was performed using the following conversions: *magnitude* (nM) points to the maximal value minus the minimal value of a series. *tmax* refers to the time point that the curve reaches the maximal value, scaled by time range (540 s). Parameter *ylast* indicates the terminal value, scaled according to the magnitude. *absdev* is a value indicating how much the time curve is deviating from a straight line (red line); calculated are at each time point the deviations from this line (green lines), and *absdev* is the average of these deviations.

**Figure 3.**
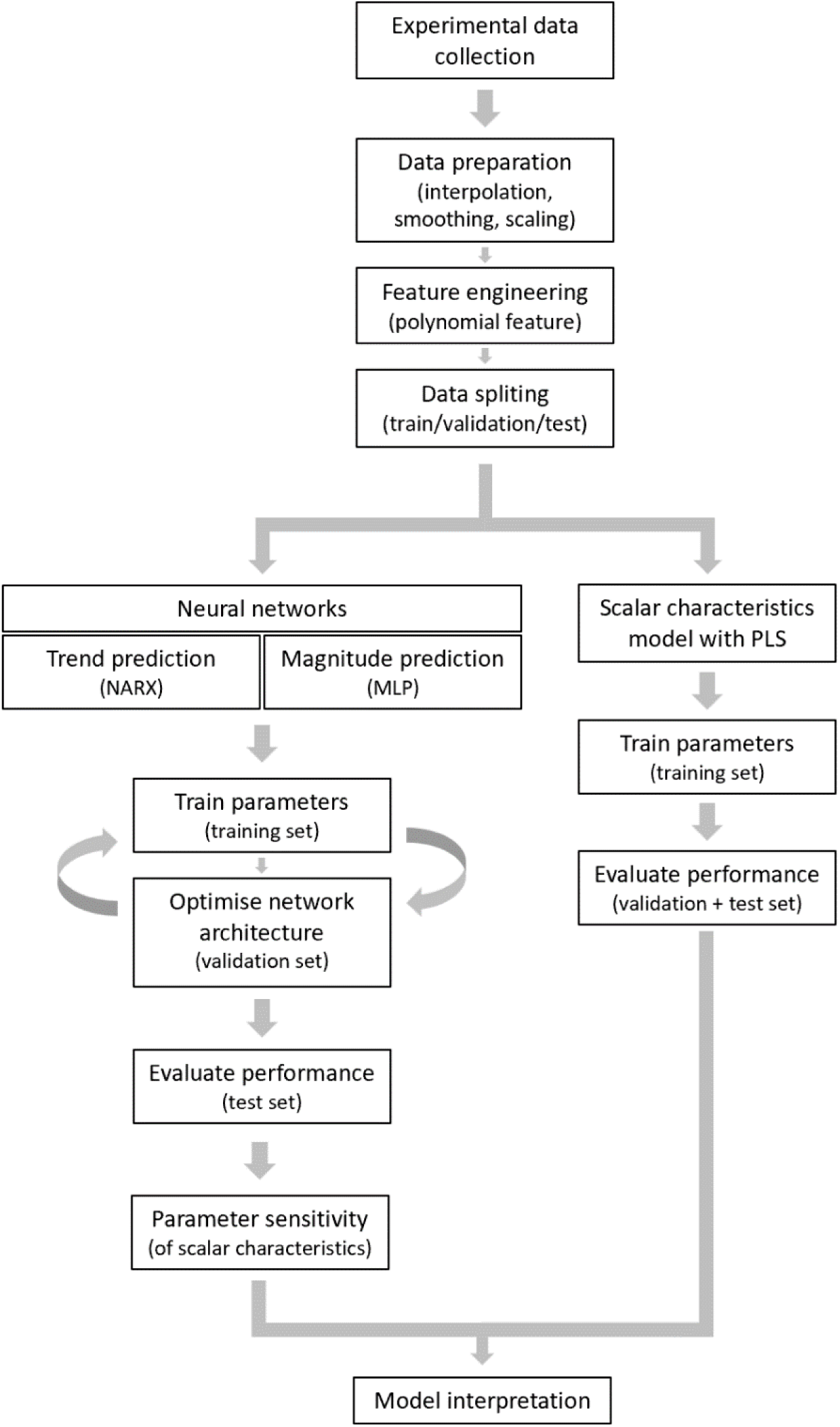
Workflow used for the data processing, neural network construction and scalar model development. For explanation, see text.

### 2.3. Neural MLP Network for Magnitude Prediction

We first aimed to better understand how the smoothened [Ca^2+^]_i_ curves of platelets relied on the various experimental conditions (CaCl_2_/EGTA, agonist dose, AI or thapsigargin). For this purpose, we generated a simple network able to predict the magnitude of the Ca^2+^ signal. The constructed multilayer perceptron (MLP) network was trained and validated, from which itappeared that the best MLP architecture had three nodes with a single hidden layer (Figure 1A). The results for the training, validation and test sets are shown in Figure S5. Plots were generated to compare the experimental data with the predictions in log scale of linear scale. Herein, each data point represents the experimental values and predicted magnitude values. These plots indicated an overall reasonable fitting, expecially for the log-scale setting.

The obtained MLP parameters associated with each node are shown in Figure S6, with a colored way of the relative weights of the combined and squared parameters in the network. A limitation of this MLP approach is that only the curve magnitude is predicted and not the curve shape.

### 2.4. Neural NARX Network for Trend Prediction

For prediction of the shape of trend of the [Ca^2+^]_i_ curves with all different amplitudes, uniform scaling is needed. Predictions modelling on these scaled time curves were made by constructing a recurrent, closed-loop neural network (NARX). For the training of the network, we used 58 scaled curves (Figure S7), which resulted in the best results for a network architecture with 3 hidden layers and 4 x 12 x 4 nodes (mean R^2^ = 0.84) (Figure 1B). The validity of the network was overall confirmed for the validation set of 7 curves (mean R^2^ = 0.71) (Figure S8). Fitting was less for the test set with 7 curves (R^2^ = 0.64), in particular for the transient curve of Exp. 58 with thrombin (Figure 4A-G). For comparison, testing the same (unscaled) amplitude curves with the MLP network resulted in a good prediction, especially for the high-magnitude curves (Figure 4H).

**Figure 4.**
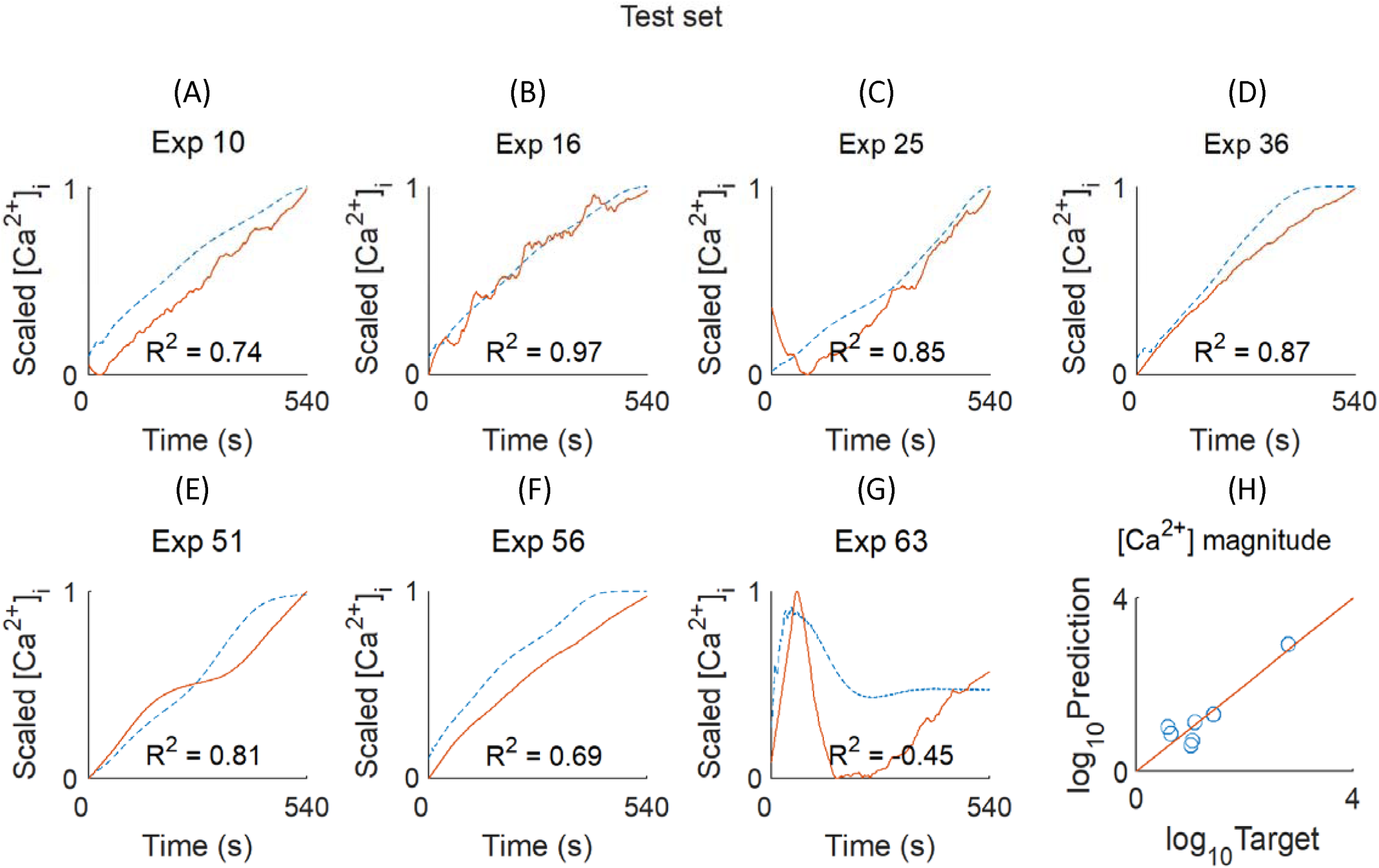
Test set results of NARX neural network to predict curve trends. (**A-G**) Testing of trend prediction of scaled [Ca^2+^]_i_ curves. The test set with experimental conditions was, as defined in Table 1. Red solid lines = actual values, blue dashed lines = predicted values. Calculated R^2^ indicated per experimental condition (negative R^2^ indicates an explained variance worse than random). (**H**) For comparison, results for the same test set are shown for curve magnitude predictions with the MLP network. Shown are the target and predicted nM levels of [Ca^2+^]_i_ in log scale.

The NARX prediction trends also provided information on the non-linear shape of the [Ca^2+^]_i_ curves. Examining the trend values of R^2^, it appeared that these were negative for Exp. 58 (Figure S8) and Exp. 63 (Figure 4). This pointed to an explained variance worse than random, and hence inability of fitting. Furthermore, also other Exp. 67, 70 and 72 with thrombin as agonist gave an R^2^ <0.4. The likely explanation with this is the transiency of the thrombin-induced [Ca^2+^]_i_ rises. The above results prompted us to compare the neural network results of both magnitude and trend prediction.

### 2.5. Combining the MLP and NARX Networks

For a combined network curve prediction, we used the training set of 58 curves (Figure S9). The training with respect to magnitude and trend predictions was then validated and tested using the remaining 14 curves (Figure S10). The combined prediction resulted in a generally improved outcome. We also performed a one-factor-at-a-time (OAT) analysis by varying the agonist concentration at different inhibitor combinations, as shown in Figure 5A (as scaled variant curves) and Figure 5B (as scalar heatmaps). For additional visualization, also the unscaled curves are represented in Figure S11, which shows both the size and trend changes of the curves.

**Figure 5.**
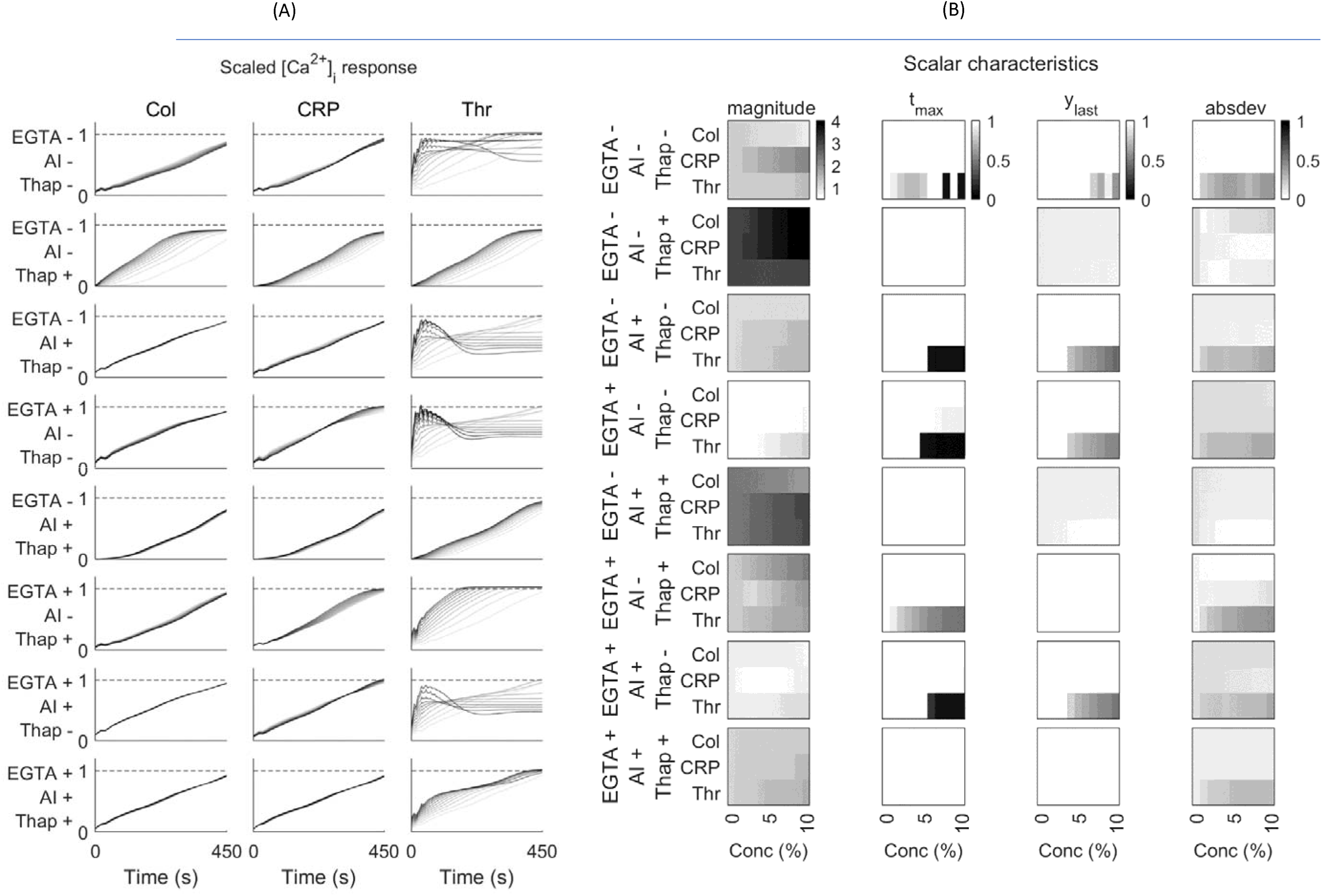
Combined variation of trend prediction of scaled platelet [Ca^2+^]_i_ responses at varying agonist concentrations. (**A**) Panels indicate prediction efficacy per agonist concentration. Lightest grey lines represent basal levels, while darker lines point to curve predictions in the presence of agonist by 1% from the basal level to 10% of the maximum concentration in the training set. Columns show conditions with indicated agonists (collagen, Col), CRP or thrombin (Thr). Rows represent different inhibitor conditions: + or - mean presence or not; from top to bottom: EGTA, apyrase plus indomethacin (AI), and thapsigargin (Thap). Dash lines at scale 1 indicate maximal trend per experiment. (**B**) Sensitivity of scalar characteristics of time curves generated by MLP and NARX model. Columns here indicate: [Ca^2+^]_i_ level at log10 base (*magnitude*), time of [Ca^2+^]_i_ (*tmax*), final [Ca^2+^]_i_ level (*ylast*), and mean absolute deviation from linear (*absdev*). All data shown are scaled at 0-1. For the unscaled [Ca^2+^]_i_ data, see Figure S7.

The OAT sensitivity analysis of Figure 5A shows how the predicted trend changed with the experimental input condition. It appeared that the predicted *magnitude* of the ‘no inhibitor’ (0 0 0) condition was mostly changed with the CRP concentration, when compared to collagen or thrombin. The presence of EGTA reduced the overall *magnitude* with to all agonists. Thrombin affected the *magnitude* most, while collagen and CRP had smaller effects. Furthermore, the presence of thapsigargin increased the overall *magnitude* prediction, regardless of the type of agonist. Furthermore, the predicted *magnitude* increased less with the concentration of thrombin, than that of collagen or CRP.

As shown in the heatmap in Figure 5B, we also compared the three scalar characteristics (*tmax*, *ylast* , and *absdev*) of the scaled curves (see Figure 2). The [Ca^2+^]_i_ peak time (*tmax*) provided information on the carve transiency. If the *tmax* was equal to the final time point (540 s) usually indicated means that the Ca^2+^ response increased over the time range. In particular with thrombin the *tmax* was often <540 s, indicating a peaking and transient response. With CRP this was only seen to a limited extent at some inhibitor conditions.

The final level of [Ca^2+^]_i_, i.e. the parameter *ylast*, displayed similar trends as *tmax* (Figure 5B). In general, *ylast* is close to 1 under conditions of a continuous increase in [Ca^2+^]_i_, and <1 when the trend hit a peak before decreasing. Thus, thrombin inducing a non-linear curve pattern produced lower *ylast* values, even at higher agonist concentrations. The parameter *absdev* (absolute deviation from a straight line) indicated how the curve deviates from a linear response. Analysis of *absdev* showed that most of curves with thrombin were non-monotonic, except for conditions at which both thapsigargin and AI were present, i.e. resulting in more linear curves (Figure 5B). Accordingly, the three scaled curve characteristics provided additional information on the Ca^2+^ response patterns upon varying the CaCl_2_, thapsigargin, AI and agonist concentrations.

From combining the results of the two tested MLP and NARX networks, several conclusions can be drawn. The transient [Ca^2+^]_i_ responses with thrombin were harder to model than the non-transient responses with other agonists. For both the weak GPVI agonist collagen and the strong agonist CRP, the scaling approach showed a mostly monotonic curve increase, being close to linear at low agonist concentrations. Furthermore, the combined magnitude and trend modelling indicated for CRP additive effects of the absence of Ca^2+^ entry (EGTA, Exp. 36), absence of secondary mediators (AI, Exp. 42), of which the former was stronger (Exp. 44). However, in spite of these insights, the black-box nature of neural network approaches could have hidden other relevant relations between curves.

### 2.6. PLS Regression Analysis

As a more straightforward approach, we also directly investigated the contribution of each experimental variable (agonist dose, EGTA/CaCl_2_, IA, thapsigargin) to the scalar curve characteristic, i.e. reducing the [Ca^2+^]_i_ time curves to *tmax*, *ylast* and *absdev*. For that purpose, we used a PLS regression analysis to fit the relationships. As the PLS regression is a linear model, it is easier to investigate the impact of input variables on the output.

As input for the PLS model, we normalised all experimental conditions to separate variables of the concentrations of agonist and inhibitors (collagen dose, CRP dose, thrombin dose, EGTA/CaCl_2_, AI, thapsigargin), all varying from 0 (none) to 1 (maximum). This resulted in a six-component model explaining the variance per component. As indicated in Figure S12, only the first two components contributed to the variance of the target. Accordingly, we fitted the 2-component PLS regression for curve *magnitude*, *tmax*, *ylast* and *absdev*, using the same training of 58 experimental conditions, while keeping the remaining 14 (previously validation and test sets) as test set of the PLS model. The variable loading coefficients of each PLS component are shown in Figure 6. This type of regression analysis was then used to predict the scalar characteristics of the test set. It also generated regression errors of both the training and test sets, which provided information on overfitting (Figure S13).

**Figure 6.**
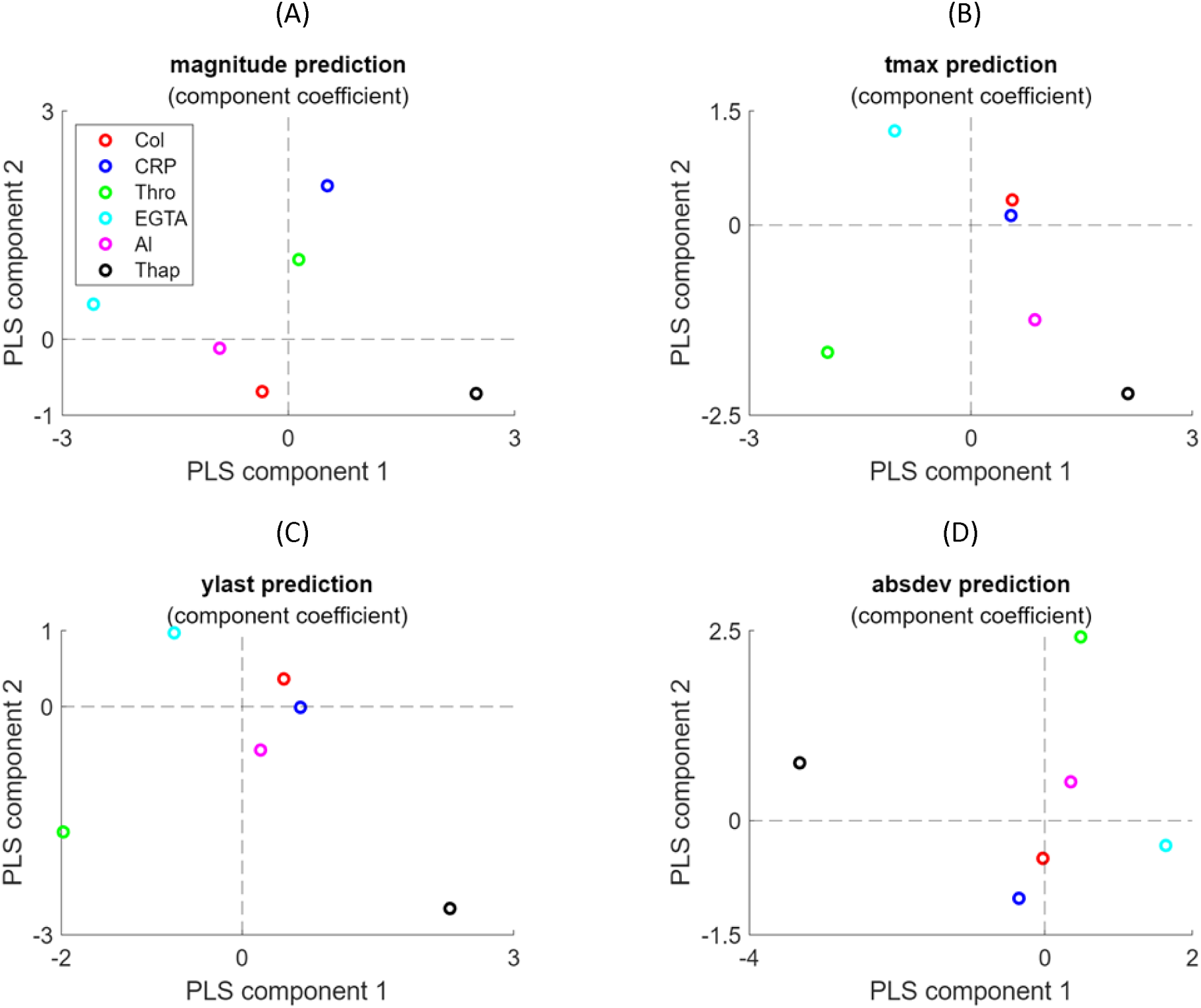
Loading coefficients of experimental variables in PLS regression analysis. Plots show for the first2 principal components the loading of the six experimental variables (collagen dose, CRP dose, thrombin dose, EGTA/CaCl_2_, AI, thapsigargin). The PLS regression analysis was performed for prediction of curve *magnitude* (**A**), *tmax* (**B**), *ylast* (**C**) and *absdev* (**D**). Indicated in colours are the contributions per varible.

In agreement with the analysis above, it appeared that the first PLS component in the *magnitude* prediction had a negative loading in the presence of EGTA and/or AI, indicating a lower level of [Ca^2+^]_i_ (Figure 6A). On the other hand, the presence of thapsigargin resulted in a highly positive loading, due to an increased [Ca^2+^]_i_ level. Indeed, the presence of EGTA stopped the entry of extracellular Ca^2+^, whereas thapsigargin increased this process by inhibiting the SERCA-type Ca^2+^ pumps controlling the STIM1-Orai1 entry pathway^17^. Furthermore, the *tlast* and *y m a x*predictions showed an opposite loading in component 1 for thrombin (negative) and thapsigargin (positive) (Figure 6B-C). This reflected the more transient, non-linear Ca^2+^ responses with thrombin in comparison to those with thapsigargin. Regarding the *absdev* prediction, the thrombin condition showed a particularly high positive weight in component 2 (Figure 6D). The NARX model showed that the patterns of Ca^2+^ curves modelled were accurately predicted, while the PLS model provided valuable understandable data regarding the significance of each variable.

## 3. Discussion

The combination of modelling approaches presented in this work introduces a new way to predict the response pattern of agonist-induced platelet Ca^2+^ responses under a great variety of conditions. The constructed neural networks by MLP and NARX were able to produce mostly correct magnitude curves of [Ca^2+^]_i_, whereas the modelling by PLS regression captured the characteristic curve shape. Our work thereby adds to the idea of a platelet Ca^2+^ calculator introduced by Diamond and colleagues [14], in that also curve patterns can be predicted by without mathematical modelling. On the other hand, we did not consider the synergistic effects of agonist combinations such as presented in that study.

It is important to note that while the present machine-learning techniques were able to fit most of the input data, the obtained output does not give a direct biological interpretation, although sensitivity analysis was used to improve the interpretability. This contrast to other modelling approaches with a clear biological meaning, such as enzyme or receptor reaction rates in ODE-based kinetic models. However, the latter approaches cannot easily capture interactions between individual signaling steps, for instance due to combinations of agonists and inhibitors.

Both the NARX network and the PLS regression analysis yielded useful results for the [Ca^2+^]_i_ curve analysis. Thus, the up to magnitude differences between traces in the presence of CaCl_2_ or EGTA and CaCl_2_ (caused by Ca^2+^ entry into the platelets) were captured by both the MLP and PLS regression models. The prediction results - i.e. sensitivity analysis for MLP and PLS component analysis for PLS -were well interpretable for this case. On the other hand, NARX could better then PLS capture the curve effects by certain experimental variables. The curve magnitude and other characteristic effects (*tmax* and *absdev*) caused by thapsigargin, was also captured by NARX, but not by PLS regression. This illustrates that neural networks as NARX can easily handle non-linear effects function due to the complex activation functions, while PLS relies in linear regression analysis and hence cannot adapt to non-linearity.

A specific problem encountered was the different shapes of the [Ca^2+^]_i_ curves used for training by the various approaches, *i.e*. more often transient with thrombin and usually linear with CRP or collagen. Although neural networks can capture any function, they need many data to train for such curve differences. In our case, only a limited number curves with either agonist could be used for training, which caused a certain imbalance in the training set. One way to fix this problem is to use data augmentation, for example by the synthetic minority oversampling technique (SMOTE) [16], which is more often used for imbalanced datasets.

In the present paper, we trained all models using platelets derived from a single donor stimulated with a range of agonists and inhibitors, which thus resulted in a new tool for investigating the complex Ca^2+^ signaling pathways in single donor platelet activation. The models can now be used to generate hypotheses for additional experimentation and to provide insights that are otherwise not obtained by traditional analytical approaches. However, appropriate use of the models is important, ensuring that the data used for training are representative, while independent data are available for validation. The use of blood from a single donor can be seen as a limitation of the study, also because this reduced the number of variable experimental conditions and, accordingly, the machine learning models had a limited predictive power. Comparing the platelet responses from multiple donors will increase the number of samples available for model building, and may thereby decrease the accuracy of the predictions for each donor.

A solution to this issue is the approach of transfer learning [17], in which a generic model can be built for the samples from various donors, and then refine the model to obtain adjusted the weights for each donor separately. This approach is being used to build personalized models for drug development [18]. An alternative is to train an auto-encoder in learning from a reduced part of the input, this to reproduce the output; this will also allow training on the data from several blood donors. Regardless of the approach followed, modelled analysis will be interesting of the effects of additional inhibitors of relevant Ca^2+^ signaling pathways, such as P2X_1_ Ca^2+^ channel antagonists [19] or STIM1-Orai1 pathways blockers [20].

Differently from the neural network models, the PLS regression analysis performed better with the relatively small sample size from one blood donor. The PLS approach is also less prone to overfitting. The present PLS regression analysis to predict the (scaled) [Ca^2+^]_i_ curve features easily allows for comparisons with the platelets from more donors. In work of the Diamond laboratory [14], a NARX model was generalized by fitting multiple networks constructed from several donors, and the determining their average prediction. Our analysis indicates that this can be done more easily by PLS regression approaches.

## 4. Methodology

### 4.1. Materials

Human α-thrombin came from Kordia (Leiden, The Netherlands); cross-linked collagen-related peptide (CRP-XL) from the University of Cambridge (UK); Fura-2 acetoxymethyl ester from Invitrogen (Carlsbad CA, USA); Pluronic F-127 from Molecular Probes (Eugene OR, USA). Horm-type collagen was obtained from Nycomed (Hoofddorp, The Netherlands). Other materials were from sources described before [21].

### 4.2. Blood Collection and Platelet Preparation

The study was approved by the Medical Ethics Committee of Maastricht University. Blood donor age and sex could not be recorded. Blood was taken into 3.2% sodium citrate (Vacuette tubes, Greiner Bio-One, Alphen a/d Rijn, The Netherlands) from consenting healthy volunteers who had not taken anti-platelet medication in the previous ten days. Platelet counts were within the reference range.

Platelet-rich plasma (PRP) was obtained from the citrated blood by centrifuging, after which the collected platelets were washed in the presence of apyrase (1 unit/mL), and then loaded with Fura-2 acetoxymethyl ester (3 µM) and Pluronic (0.4 µg/mL) at a count of 2 × 10^8^/mL for 40 min at room temperature, such as described before [22]. The cells were finally resuspended at a concentration of 2 × 10^8^/mL in Hepes buffer pH 7.45 (10 mM Hepes, 136 mM NaCl, 2.7 mM KCl, 2 mM MgCl_2_, 5.5 mM glucose, and 0.1% bovine serum albumin).

### 4.3. Calibrated Cytosolic Ca ^2+^ Measurements

Using Fura-2-loaded platelets, changes in cytosolic [Ca^2+^]_i_ were measured in 96-well plates with a FlexStation 3 (Molecular Devices, San Jose, CA, USA), as described [22]. When appropriate, the platelets in wells were pretreated with apyrase (0.1 unit/mL) plus indomethacin (20 µM), or with thapsigargin (1 µM) for 10 min. After the addition of 0.1 mM EGTA or 1 mM CaCl_2_, platelets were stimulated by automated pipetting with one of the following agonists: CRP (1 or 10 µg/mL), collagen (1, 3, 10 or 30 µg/mL), thrombin (0.3, 1, 3 or 10 nM), or none of these (control). In all wells, changes in Fura-2 fluorescence were measured quasi-simultaneously over time at 37°C by ratiometric fluorometry, also including appropriate control wells for calculating nM concentrations of [Ca^2+^]_i_ [22].

### 4.4. Selection of Platelet [Ca ^2+^]_i_ traces for Modelling

Calibrated agonist-induced time series of [Ca^2+^]_i_ with the various experimental conditions were performed with Fura-2-loaded platelets from 6 donors [15]. For the present modelling approach, a complete set of 72 time curves (Table 1) was chosen from one donor, which were representative for those all six donors. In the table, the validation and test conditions are highlighted in blue and red, respectively. The criteria for splitting these are indicated below.

### 4.5. Preparation of Input Data

The traces of nM changes in [Ca^2+^]_i_ in platelets for experiments involving CRP or collagen were measured every 4 s, while those for experiments with thrombin had an interval of 2-4 s. To be able to compare all 72 traces, all raw data (Figure S2) were linearly resampled and interpolated to generate 1 s time steps, from 0 s to 540 s (9 min). To minimize the noise in the dataset, the curves were smoothened with a Savitzky–Golay filter (Figure S3).

In cases where scaling of data was needed, a subset of interpolated curves was subjected to a standard min-max scaling algorithm to obtain values between 0 and 1. For the scaling of input conditions, experimental variables were set to have values in the range [-1, 1] (Table 1). Herein, -1 indicated no agonist or inhibitor present, while 1 indicated that the concentration of agonist or inhibitor was maximal across the samples.

For constructing a multilayer perceptron (MLP) network, a regression model was built using magnitudes of the [Ca^2+^]_i_ time series. The experimental variables were used as input values (Figure 1A). Herein we used the mean square error (MSE) as a cost function. This ensured a better fit for larger values (in the order of magnitude). To handle this complexity, we set the target (output) for the model as log-scaled values of the nM [Ca^2+^]_i_ range as log_10_(max – min). This optimized the overall accuracy across log scales.

Considering that the number of features was small with 6 experimental conditions (Table 1), we also generated polynomial features (quadratic feature combinations) to increase the feature number from 6 to 27. The MLP network was optimized by setting the number of hidden layers to 1, with the number of nodes randomly selected from 1 to 10. Given the relatively small training set that was available, network architecture options were chosen to train only a low number of parameters, thus preventing overfitting. Networks were trained 100 times, starting from random weights. The best structure was chosen as the one with a minimal score in the cost function of the validation set. Network training was performed using the Levenberg-Marquardt algorithm, containing a rectified linear unit (ReLU) as an activation function in each node. Modelling was conducted using Matlab R2022a and the Neural Network Toolbox.

### 4.6. Trend prediction of NARX network

Another type of neural network was constructed to predict the trends of smoothened and scaled [Ca^2+^]_i_ time curves. To better capture the time dynamics (*i.e.* the shape of the curve), we choose a non-linear autoregressive network with exogenous inputs (NARX) and parallel architecture [23,24], which is known as a closed-loop neural network. For this NARX network, the model’s output *y(t)* was used to fit the target (i.e., the smoothened and scaled [Ca^2+^]_i_ curves). The output then generated feedback as additional input to the network, when combined with the experimental condition (Figure 1B). The mathematical expression of [Ca^2+^]*(t)* is written as follows:

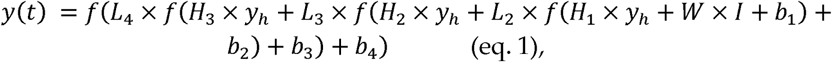

where *y(t)* is [Ca^2+^]_i_(t), *I* is an input matrix of experimental conditions, y_h_ is the feedback delay (history) of *y*. Furthermore, *W* and *H* _n_ are the input matrix’s weight and feedback delay of y, respectively; *b*_n_ are biases, and *L* _n_ are weights of each hidden layer; *f* is the activation (transfer) function. Note that the product of the matrix also is a matrix, meaning that the equation represents a summation of numerous parameters and functions.

For the feedback delays, we choose the values at the last 1, 3, 6, 10, 15, 21, 28, and 36 s prior to the current value of the [Ca^2+^]_i_ time series. These feedback delays hence kept the information about recent values, while preserving the long-term memory of the system. Initial values of the feedback delays were set to zero, as the system was assumed to be in a steady state prior to the agonist-induced activation of platelets. The use of MSE as a cost function allowed us to make predictions of the scaled min-max [Ca^2+^]_i_ time series. The scaling was performed per time series, implying that each series had the same range [0,1]. Polynomial features were also used in this network, thus expanding the number of inputs from 6 to 27.

The neural network architecture was optimized in a way to maximize the goodness of fit, but to prevent overfitting. We used three hidden layers, with each layer’s size varying between 2 to 20 nodes (not including feedback delays). This gave approximately 7000 different architectures being trained. A randomized grid search was employed to find the best architecture. For training, the Levenberg-Marquardt algorithm was used with a hyperbolic tangent sigmoid (*tansig*) as an activation function. Since parameter fitting in the neural network depended on a random seed, each architecture was fitted 100 times, after which the best parameters were used for comparison. The networks were again built and trained in Matlab.

### 4.7. Parameter Sensitivity Analysis

To perform sensitivity analysis, the method of one-factor-at-a-time (OAT) was applied [25]. This procedure keeps all other variables fixed to the central or baseline values while changing one variable at a time. Since all effects were computed with reference to the same central point in space, this improved comparability of the outcomes. As a default, we set the conditions of EGTA or CaCl_2_, autocrine inhibitors (AI) or not, and thapsigargin or not as 1 or 0 (2^3^ = 8 possible combinations). Furthermore, the agonist concentration was scaled from 0 to 10% of the maximal concentration, i.e. 30 µg/mL collagen, 10 µg/mL CRP or 10 nM thrombin. The shape of each [Ca^2+^]_i_ time curves was defined according to their scalar characteristics, namely the magnitude of the response, the peak time, the relative terminal level, and the mean deviation from a straight line, such as indicated in Figure 2.

### 4.8. Partial Least Square Regression Analysis

PLS regression analysis [26,27] was used as an extension of principal component analysis, in which PLS instead of maximizing the variance in each component, maximizes the covariance between an input matrix X and an output matrix Y. Herein, each component has a latent variable *t*_i_, while the linearly weighted combination of the latent variables generates the prediction of outcomes (Y matrix) as follows:

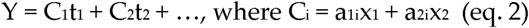

The experimental conditions of Table 1 were used as the X matrix, and the scalar characteristics of each [Ca^2+^]_i_ time series served as Y matrix. The number of components in the PLS analysis was obtained from the optimal variance achieved, when increasing the components. The loading weights depended on input variables that were most important for the predictions. By maximizing the covariance between the explanatory variable X and the response variable Y, the most relevant components in X were obtained that affected the changes in Y. Stated otherwise, by examining the loading weights of first few latent variables that accounted for the majority of explained covariance, we could identify those experimental conditions with a most significant impact on [Ca^2+^]_i_ time curves.

## Author Contributions

Conceptualization, methodology and formal analysis: C.T., J.L.D. and R.C. Investigation: C.T. and H.Y.F.C. Resources and supervision: J.M.G., J.W.M.H. and R.C. Data curation: C.T. and H.Y.F.C. Writing – original draft preparation: C.T. and R.C. Writing – review and editing: CT, J.W.M.H. and R.C. Funding acquisition: J.M.G. and J.W.M.H.

## Institutional Review Board Statement

The study was approved by the local Medical Ethics Committees (Maastricht University Medical Centre, NL31480.068.10). All subjects gave full informed consent according to the Declaration of Helsinki, and all methods were performed in accordance with the relevant guidelines and regulations.

## Informed Consent Statement

Informed consent was obtained from all subjects involved in the study. According to ethical per-mission, all subjects gave blood without tracing samples to certain individuals.

## Data Availability Statement

All data are included in the manuscript as figures, tables, or supplementary figures.

