## Supplemental information for "Extended Modelling of Platelet Calcium Signaling by Combined Recurrent Neural Network and Partial Least Squares Analyses"

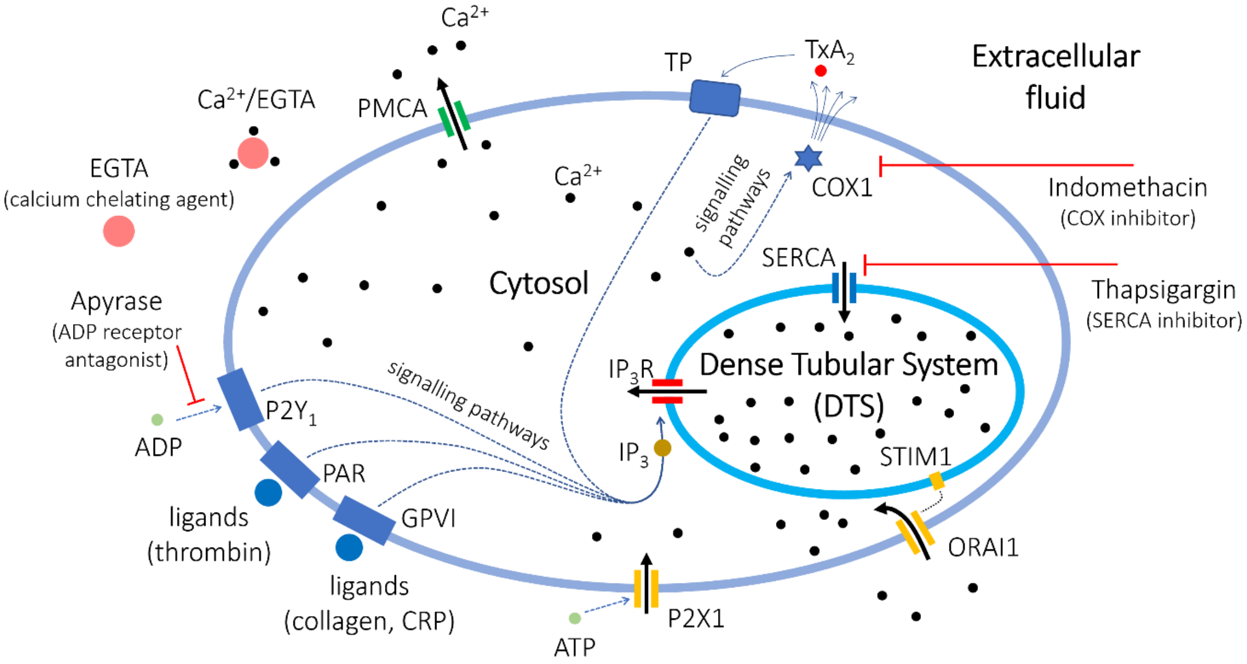


**Figure S1. Overview of receptor-induced $Ca^{2+}$ signaling mechanisms in platelets.** Collagen and collagen-related peptide (CRP) activate platelets via glycoprotein GPVI (GPVI), while thrombin acts by cleaving proteinase-activated receptors (PAR). Ligand binding to these receptors induces the formation of inositol 1,4,5-trisphosphate ($IP_3$), which stimulates the $IP_3$ receptors ($IP_3R$) in the membrane of the dense tubular system (DTS). This stimulation leads to a discharge of $Ca^{2+}$ from intracellular stores into the cytosol. Entry of extracellular $Ca^{2+}$ is mediated by the Orai1 $Ca^{2+}$ channels in the plasma membrane, which couple to STIM1 $Ca^{2+}$ sensors in the DTS membrane. A fast and quickly desensitized entry of $Ca^{2+}$ is mediated by ATP, activating the $P2X_1$ ion channels. In addition, autocrine produced ADP and $TxA_2$, via their receptors, potentiate the $IP_3$ production. Back pumping of released $Ca^{2+}$ out of the cytosol occurs by SERCA $Ca^{2+}$ ATPases in the DTS, which are inhibited by thapsigargin. Back pumping out of the cells occurs by the PMCA $Ca^{2+}$ ATPases. The presence of extracellular $CaCl_2$ or EGTA allows $Ca^{2+}$ entry or not. The formation of $TxA_2$ is inhibited by indomethacin, whereas the effects of ADP are blocked by apyrase (indicated in text as autocrine inhibitors, AI).

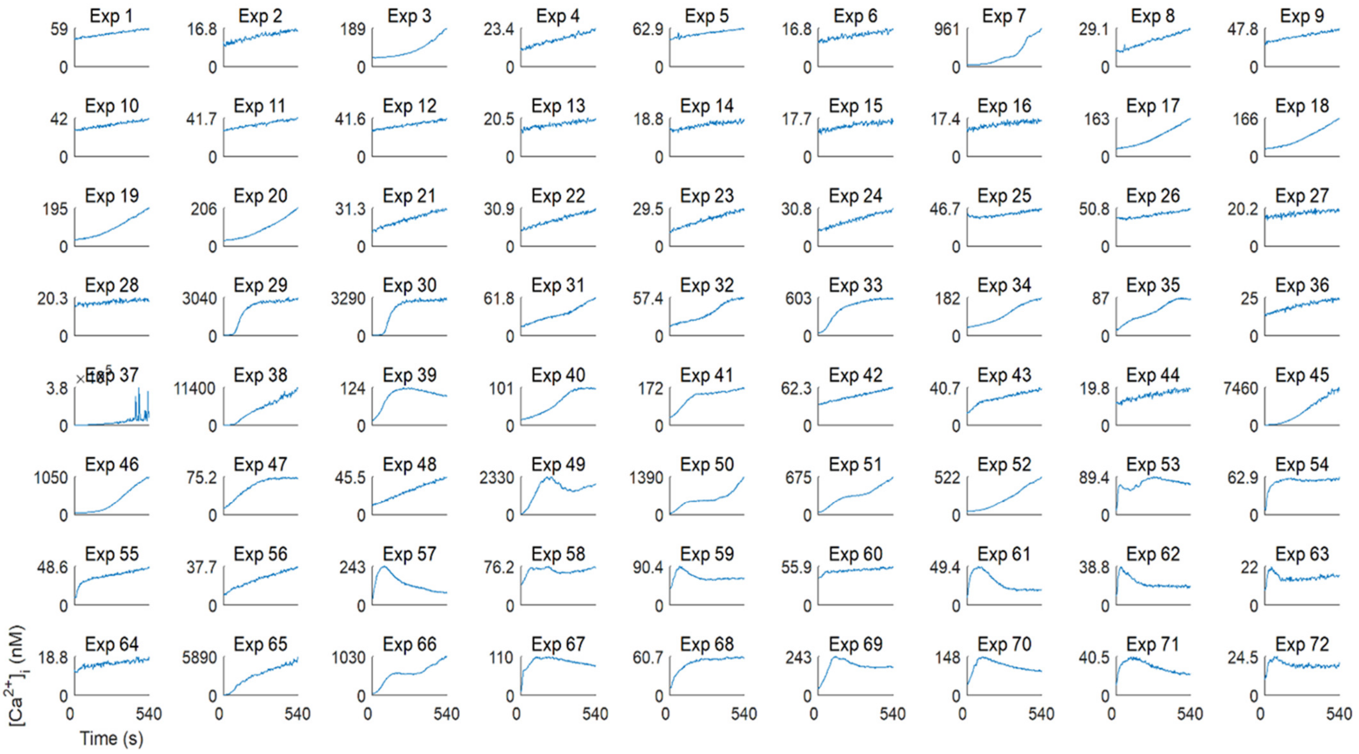


**Figure S2. Raw data of 72 agonist-induced $[Ca^{2+}]_i$ time curves of one batch of platelets used as entry for the modelling studies.** Note the widely different ranges of nanomolar levels of $[Ca^{2+}]_i$ per condition (Exp. 1-72). *Y*-axes represent linear ranges in nM $[Ca^{2+}]_i$. Time axes are set at 0 to 540 s. For the different experimental conditions, see Table 1.

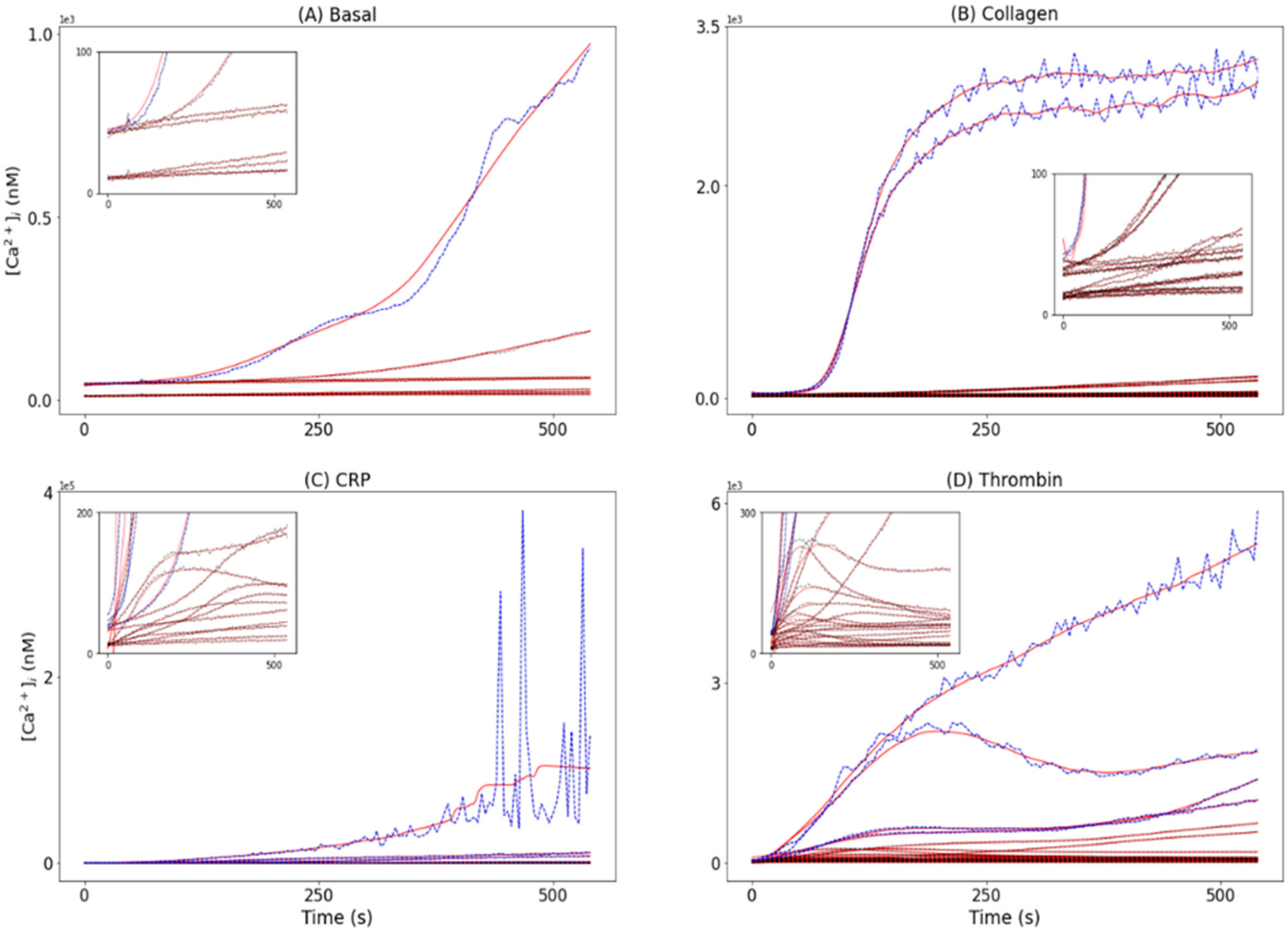


**Figure S3. Raw and smoothened agonist-induced platelet $[Ca^{2+}]_i$ time curves.** The 72 experimental conditions using Fura-2-loaded platelets (Table 1) were grouped into four panels according to the agonist used: (**A**) basal (no agonist), (**B**) collagen, (**C**) CRP, or (**D**) thrombin. Resampled and interpolated curves were smoothened with a Savitzky-Golay filter. Shown are the original curves (dash black lines) and the filtered curves (red lines). Note that high supra-micromolar rises in $[Ca^{2+}]_i$ were obtained in the presence of thapsigargin and $CaCl_2$ (dashed blue lines).

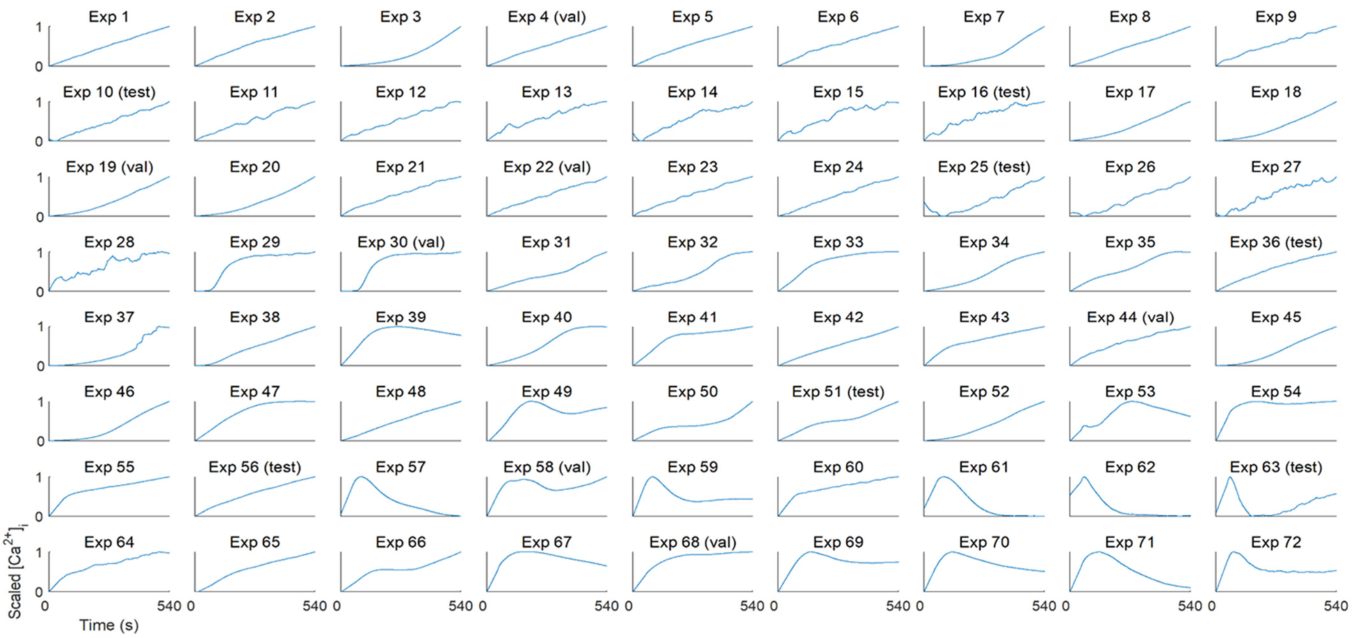


**Figure S4. Univariate scaled agonist-induced $[Ca^{2+}]_i$ time curves of platelets.** Numbers for experiments and agonist/treatment conditions are indicated in Table 1. Raw input data in nM were curve interpolated, smoothened and linearly scaled 0-1. Time axes are from 0 to 540 s. For the different experimenta conditions (Exp. 1-72), see Table 1.

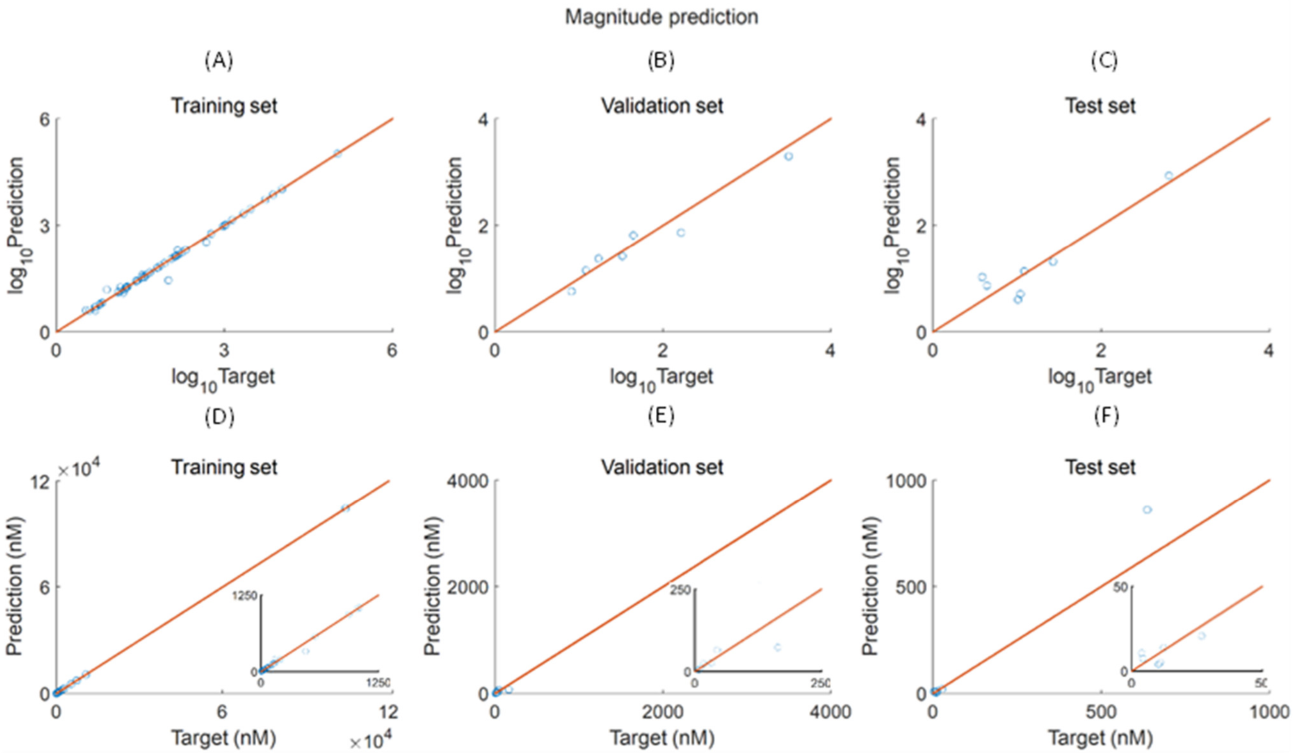


**Figure S5. Magnitude prediction of agonist-induced platelet [$Ca^{2+}$]$_i$ time curves**. Shown are for the time curves of the selected training set (**A, D**), validation set (**B, E**) and test set (**C, F**), the relation between measured target levels and predicted levels (in nM). Y-axes are in log10 scale (**A-C**), or in linear scale (**D-F**). Curve selection for the validation set were Exp. 4, 19, 22, 30, 44, 58, 68); for the test set Expt. 10, 16, 25, 36 ,51, 56, 63; and for the training set all the rest. Red lines represent diagonals.

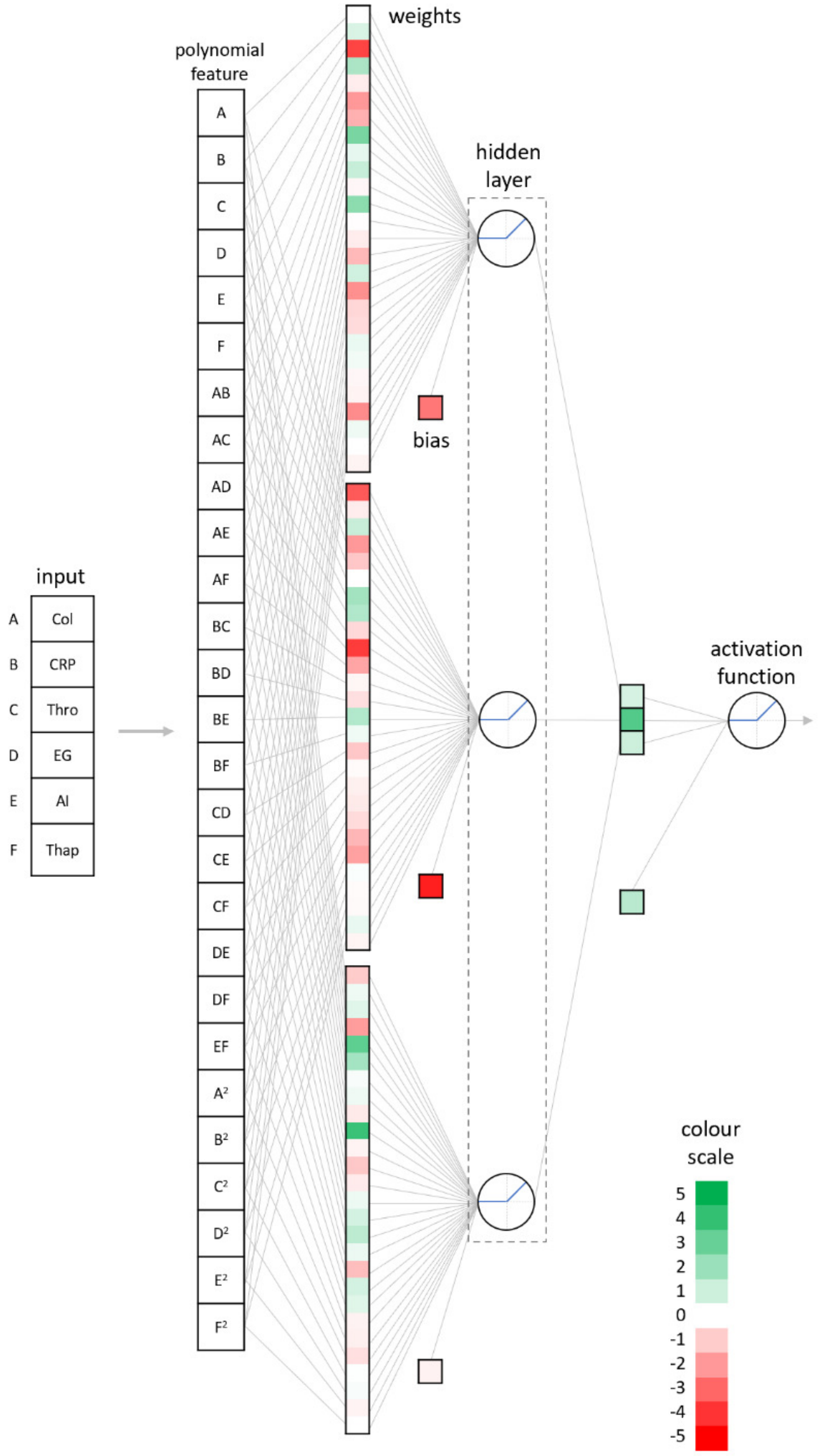


**Figure S6. Parameter composition of polynomial multilayer perceptron (MLP) network associated with each node.** Note the 27 combinations made from 6 input variables. Relative weights of per node are displayed in color scale.

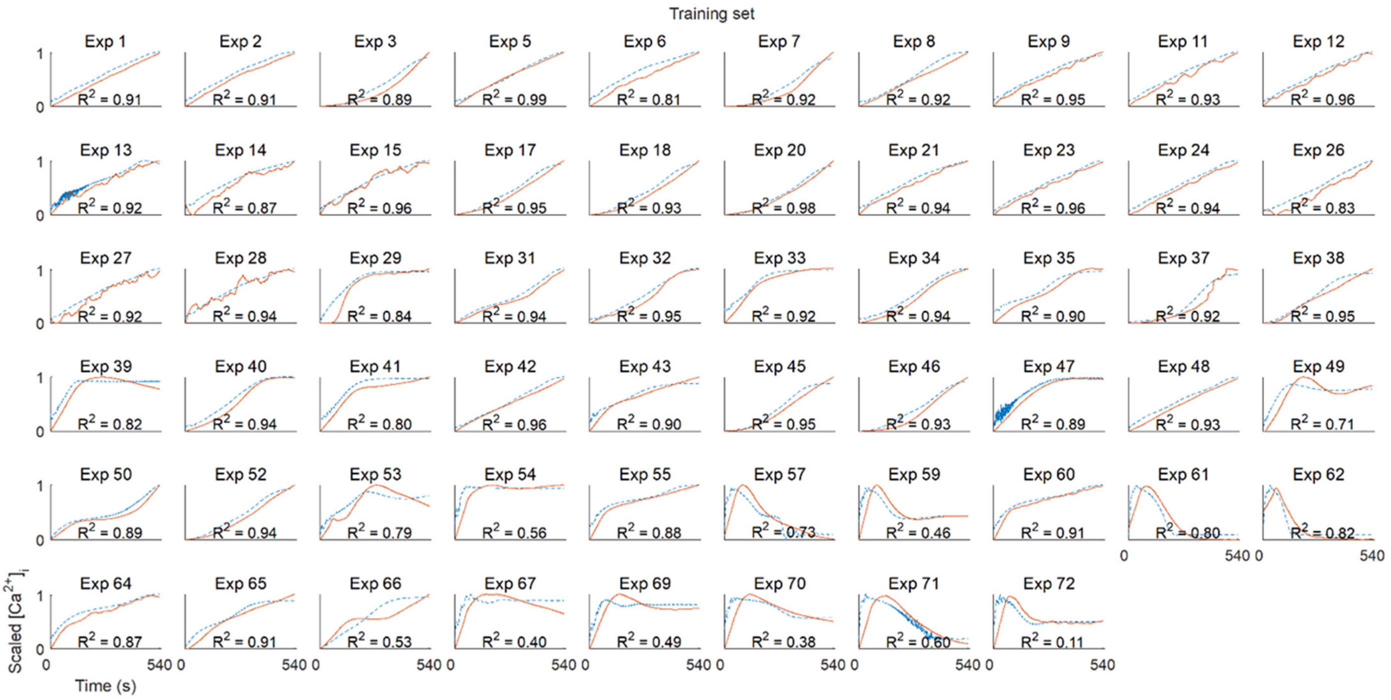


**Figure S7. Trend prediction using NARX of agonist-induced platelet $[Ca^{2+}]_i$ time curves in the training set**. Experimental numbers and agonist/treatment conditions are explained in Table 1. Vertical axes represent scaled curves from 0 to 1. Horizontal axes represent measurement time from 0 to 540 s.

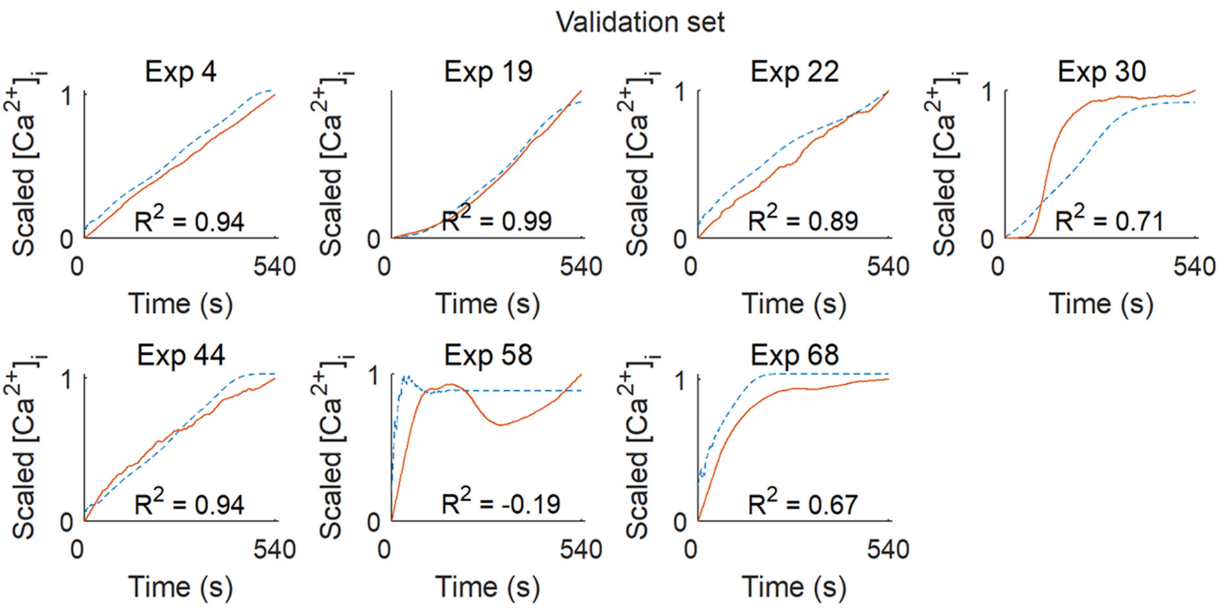


**Figure S8. Trend prediction of agonist-induced platelet $[Ca^{2+}]_i$ time curves in validation set.** Experiments were numbered as in Table 1. Vertical axes represent scaled responses from 0 to 1. Horizontal axes represent experimental time from 0 to 540 s.

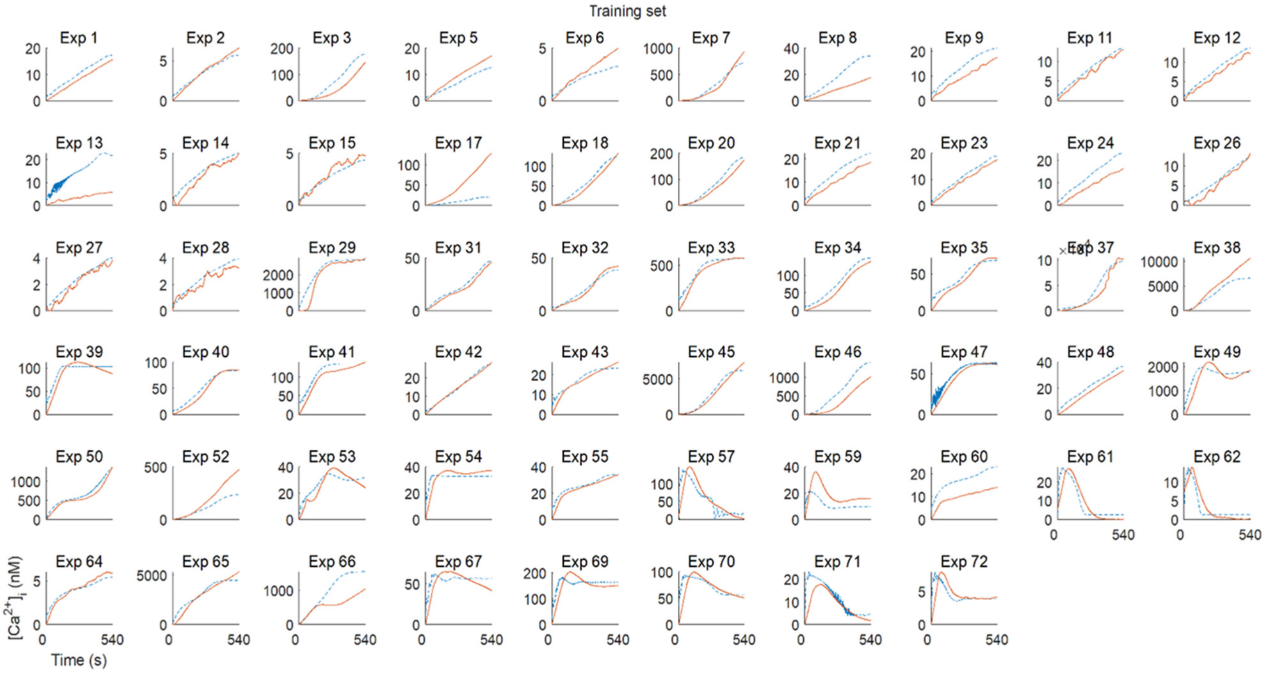


**Figure S9. Magnitude and trend prediction of platelet [$Ca^{2+}$]$_i$ time curves in training set.** The predicted curves resulted from the combination of magnitude and trend predictions. Experiments were numbered as in Table 1. Panels indicate nanomolar [$Ca^{2+}$]$_i$ levels versus time (0 to 540 s).

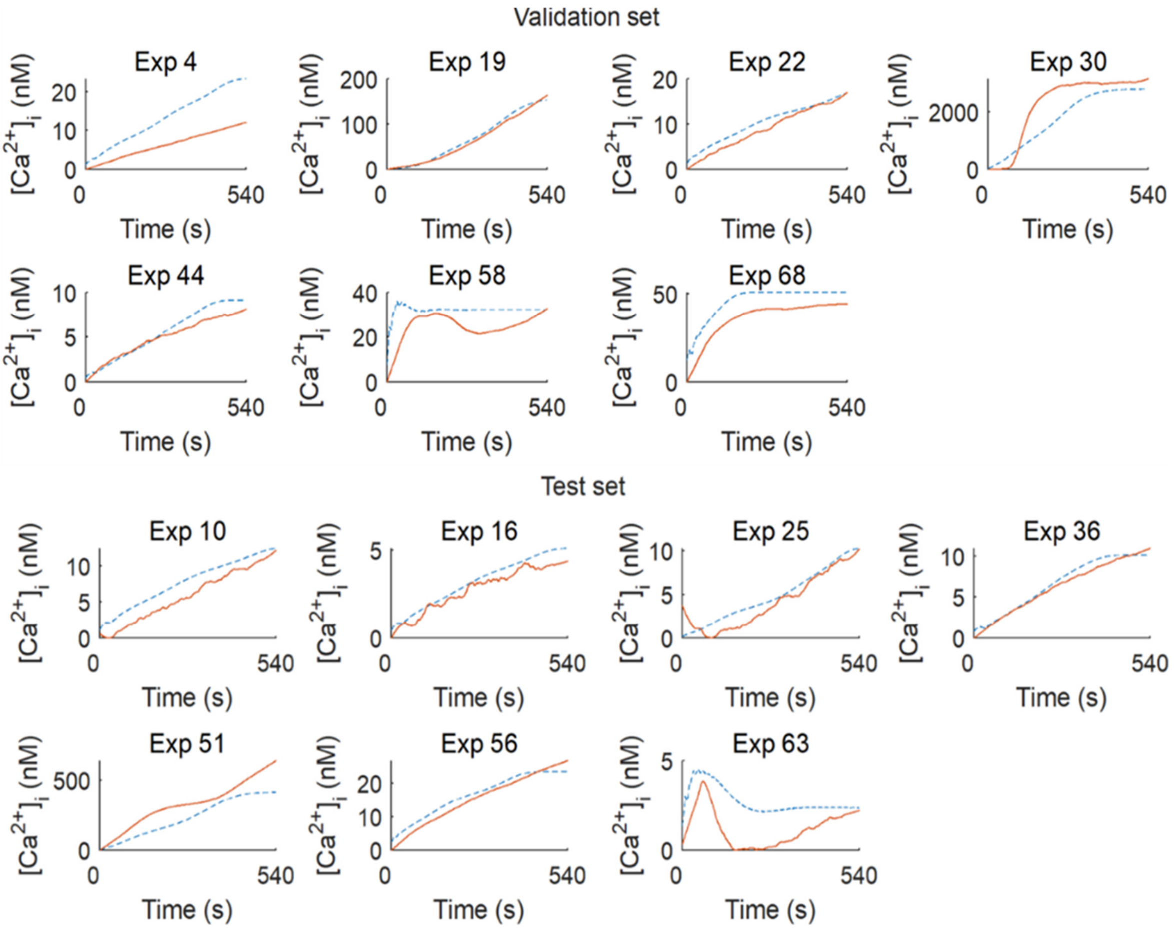


**Figure S10. Combined prediction of platelet $[Ca^{2+}]_i$ time curves with magnitude and trend predictions**. Indicated are results from the validation set (upper part) and the test set (lower part). The results from the two models were combined (magnitude and trend prediction). Experiments were numbered as in Table 1. Vertical axes indicate $[Ca^{2+}]_i$ levels in nM, and horizontal axes indicate time.

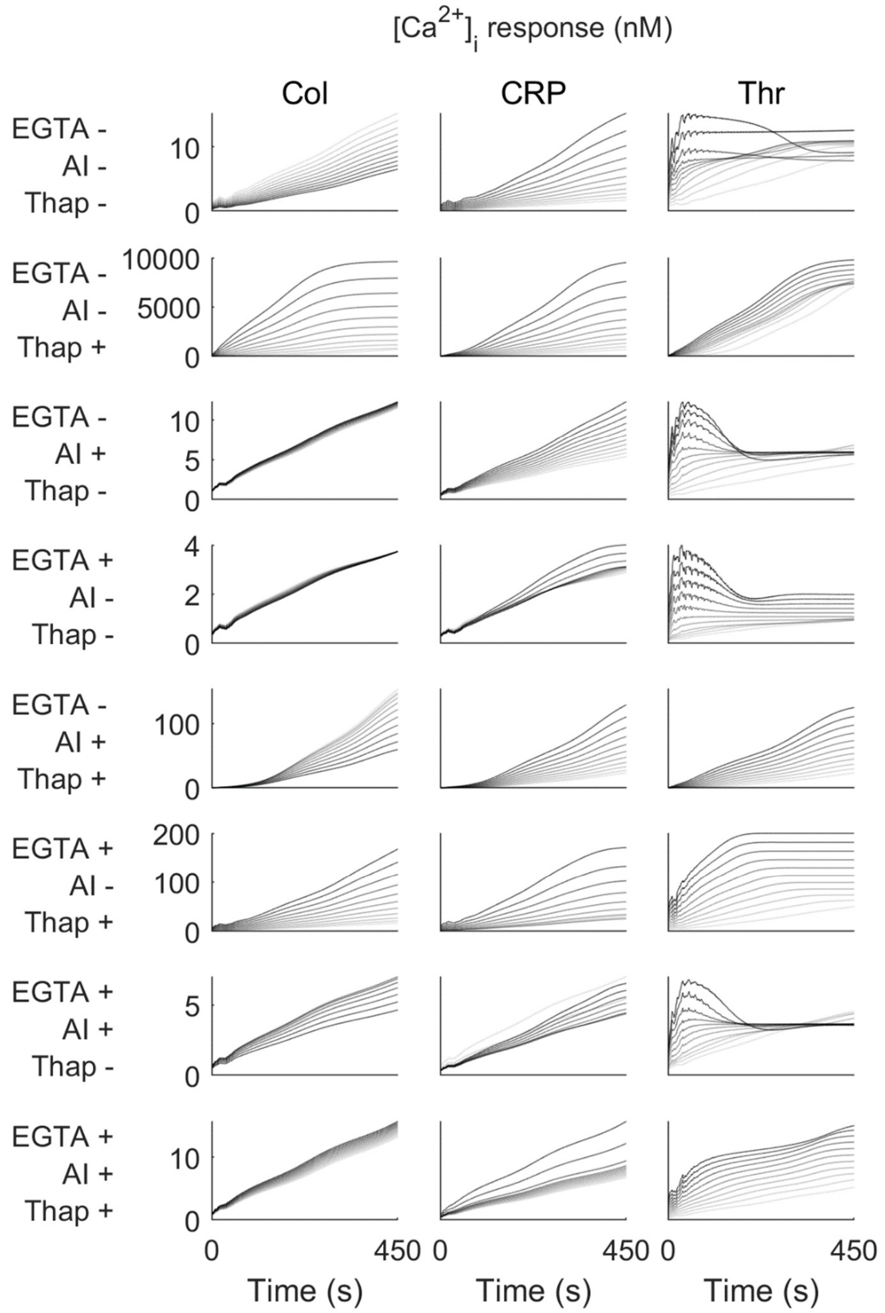


**Figure S11. Variation of trend prediction of $[Ca^{2+}]_i$ time curves with increasing agonist concentrations.** Panels indicate prediction efficacy per agonist concentration. Lightest grey lines represent basal levels, while darker lines represent a curve prediction due to an increment of ligand by 1% from basal level to 10% of the maximum concentration used in the training set. Columns show conditions with different agonists (collagen, Col), CRP or thrombin (Thr). Rows represent different inhibitor conditions: **+** or **-** indicates presence or not. From top to bottom: EGTA, apyrase plus indomethacin (AI), and thapsigargin (Thap). Shown are unscaled levels of $[Ca^{2+}]_i$ (nM); for scaled data, see Figure 5.

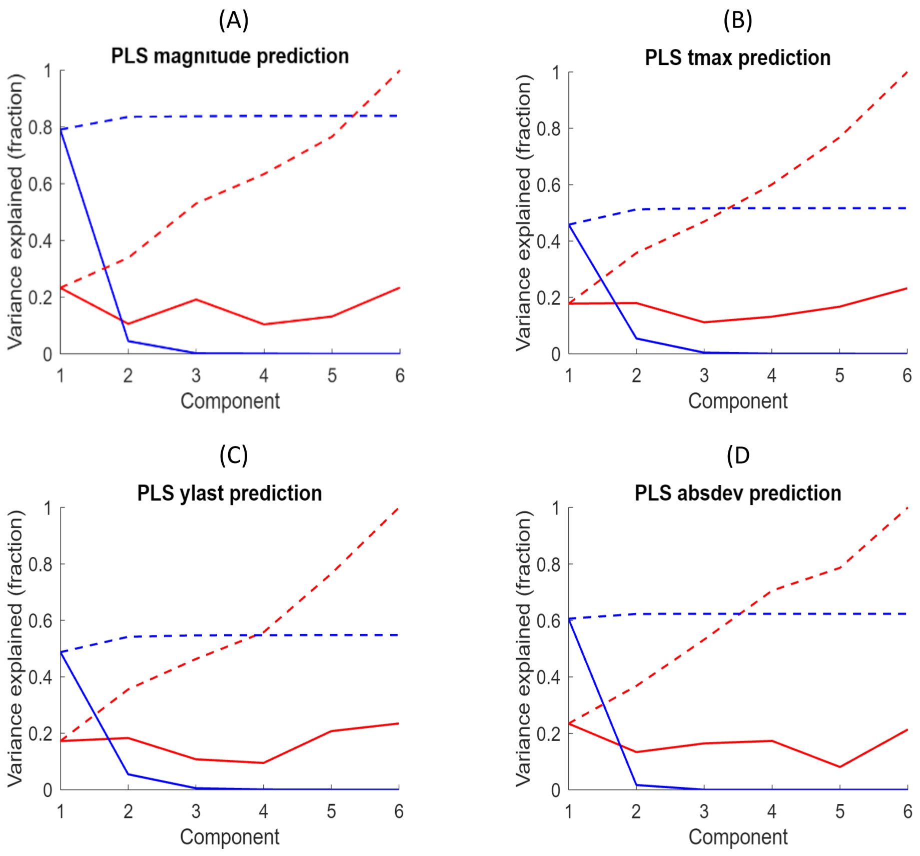


**Figure S12. Variance in PLS regression model explained per component.** Plots show per principal component of PLS, the fractions of explained variance in the dataset. A PLS regression analysis was used for the prediction of curve *magnitude* (**A**), *tmax* (**B**), *ylast* (**C**) and *absdev* (**D**). For definition of the six included experimental variables, see Figure 6. Red lines indicate the explained variance of input (experimental condition); blue lines show the explained variance of the target (curve scalar characteristic). Dashed lines display cumulative sums for increasing components. Note that only the first two PLS components contributed to the the target variance.

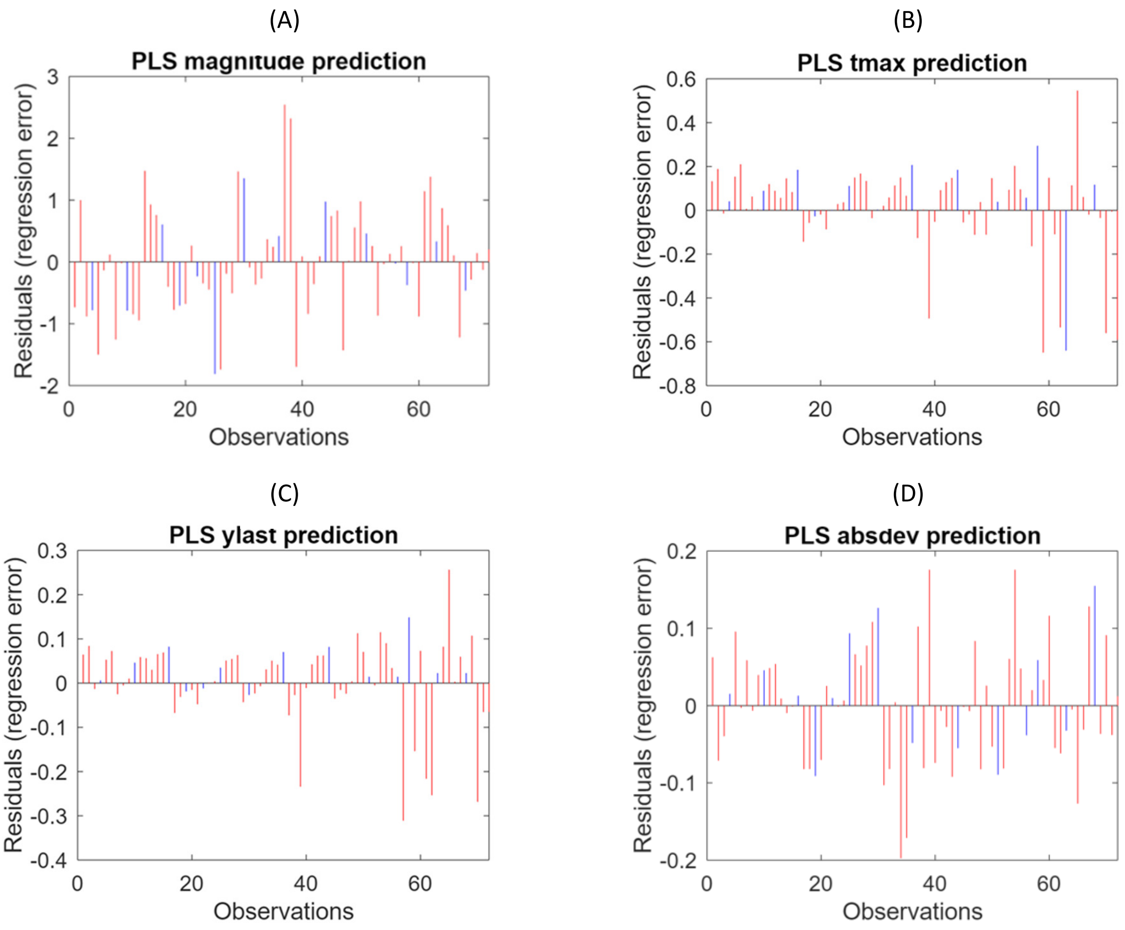


**Figure S13. Residual deviations of the PLS regression model.** Indicated are the absolute regression errors upon increasing observations in the training set (red bars) and test set (blue bars). Regression error was defined as the actual value minus the predicted value. A positive error indicates underestimation of the explained variance. PLS regression analysis was performed for prediction of the curve *magnitude* (**A**), *tmax* (**B**), *ylast* (**C**) and *absdev* (**D**).
